# Low genetic diversity in *Colobus vellerosus* populations in Kikélé Sacred and Okuta Kobunan Forests, Benin

**DOI:** 10.1101/2025.07.30.667755

**Authors:** F. Darius Accrombessi, S. S. Mireille Toyi, Inza Kone, David J. Zinner, David Djimenou, Chabi A. M. S. Djagoun, Christian Roos, Dietmar Zinner

## Abstract

The critically endangered white-thighed colobus, *Colobus vellerosus,* is on the brink of extinction, necessitating the implementation of effective conservation management strategies. The population in Kikélé village serves as the primary remaining stronghold for this species in Benin, comprising around twenty-eight individuals in the small Kikélé Sacred Forest and an additional eight individuals in the community-managed Okuta Kobunan Forest. These two populations are believed to have descended from a single founding pair introduced to the Kikélé region circa 1800. Given the small population size and the possible severe genetic bottleneck at its foundation, the genetic diversity might be extremely low. In our study, we conducted a first analysis of the genetic diversity of the two populations using mitochondrial markers, the complete cytochrome b (cytb) and a segment of the hypervariable control region (D-loop, 750 bp). Our findings revealed only one cytb haplotype, along with two haplotypes that differ by just one site in the D-loop. We recommend a range-wide population genetic assessment of the species to explore the possibility of translocations as a potential genetic rescue strategy.

## Introduction

The genus *Colobus* comprises five species of black-and-white colobus monkeys: *C. angolensis*, *C. guereza*, *C. polykomos*, *C. satanas*, and *C. vellerosus* (Fashing, 2022). Four of these species inhabit West Africa, and three of them are highly endangered. The king colobus (*Colobus polykomos*), ranging from southern Senegal to the Sassandra River in Côte d’Ivoire, is categorised as “Endangered” (Gonedelé Bi et al., 2020). The white-thighed colobus (*Colobus vellerosus*) occurs from the interfluvial area between the Sassandra and Bandama rivers in Côte d’Ivoire to western Nigeria, traversing the Dahomey Gap (Matsuda Goodwin et al., 2020). It is “Critically Endangered” (Matsuda Goodwin et al., 2020) and has been listed as one of the World’s 25 most endangered primates for several years (Mittermeier et al., 2024). The black colobus (*Colobus satanas*) occurs on Bioko (*C. s. satanas*), and from south Cameroon to Gabon (*C. s. anthracinus*) (Grubb et al., 2003) and is categorised as “Vulnerable” (Maisels & Cronin, 2020). Only the western guereza (*C. guereza occidentalis*), occurring from eastern Nigeria and Cameroon to Uganda, is categorised as “Least Concern” (Oates et al., 2020).

The range countries of the West African species have a dense and rapidly growing human population (OECD/SWAC, 2021). Forest destruction and fragmentation have been and are extensive, and the illegal hunting of wildlife is uncontrolled in most places, even in the vicinity of protected areas (CILSS, 2016; Oates, 2011; Spracklen et al., 2015). The situation is particularly dire for the white-thighed colobus, which occurs in small numbers in only a few local populations in Côte d’Ivoire, Ghana, Togo, and Benin (Schwitzer et al., 2017, 2019). Whether it still occurs in western Nigeria and Burkina Faso is questionable (Ginn & Nekaris, 2014; Oates, 2011). Currently, no protected areas in its distribution range have any in-situ conservation programs for the species, and there are no ex-situ populations (Schwitzer et al., 2017).

In Benin, Djègo-Djossou and Sinsin (2009) estimated a decrease of its range from 56,000 km^2^ to 9,000 km^2^ between 1953 and 2009, corresponding to a reduction of more than 80%. The species’ geographical range in Benin included six classified forests: Monts Kouffé, Wari-Maro, Lama Classified Forest, Pénéssoulou, Ouémé Supérieur, Ouémé Boukou, and unprotected forests as follows: Bonou Swamp Forest, Ouémé Supérieur River Swamp Forest, Lokoli Swamp Forest, Kikélé Sacred Forest (KSF) (Djègo-Djossou, 2013), and Okuta Kobunan Forest (Fig. 1). However, the species appears to have been extirpated from three of these classified forests, namely Monts Kouffé, Wari-Maro, Ouémé Supérieur, and one of the unprotected forests, the Lokoli Swamp Forest (Schwitzer et al., 2019). Recent wildlife inventory in the Pénéssoulou Forest Reserve did not find white-thighed colobus (Dossa et al., 2021). Very little is known about the species status in the Lama Classified Forest, Ouémé Supérieur, Ouémé Boukou, Bonou, and Ouémé Supérieur River Swamp Forest. The KSF is currently the only forest where a population of around 28 individuals of *C. vellerosus* can easily be observed, and where a few studies on their behavioural ecology have been carried out (Dewanou et al., 2023; Djègo-Djossou et al., 2012, 2015; Matsuda Goodwin & Chabi Ota, 2020; Ota, 2020).

**Fig. 1:**
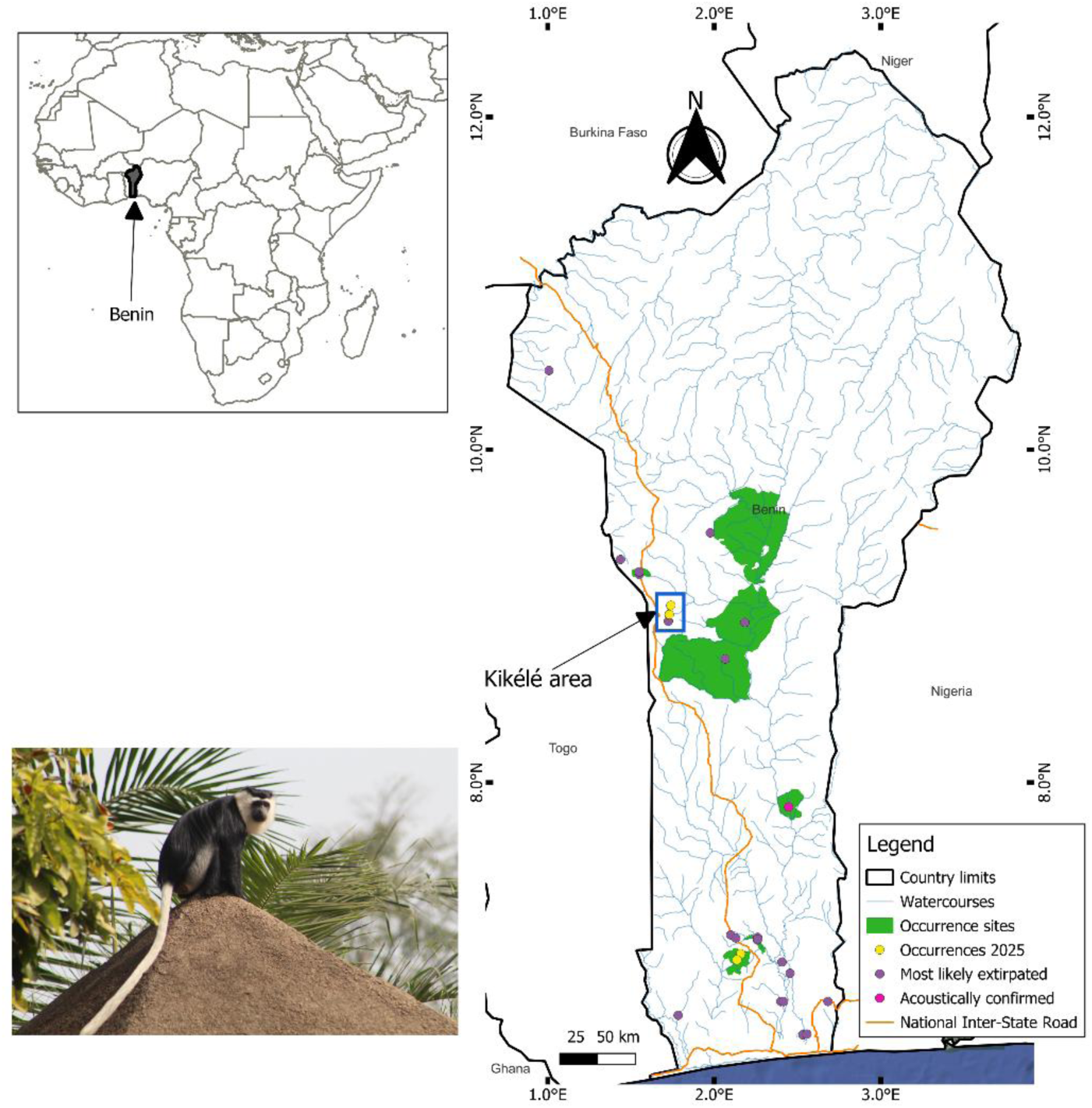
Historical and current sites of white-thighed colobus (*Colobus vellerosus*) occurrence in Benin, the Kikélé study area, and a photograph of an adult white-thighed colobus male on top of the roof of a building in Kikélé village.

The sacred character of the KSF is rooted in the totemisation of white-thighed colobus (Djègo-Djossou et al., 2012; Ota, 2020). The human residents of the Kikélé area attribute this significance to three main criteria: the belief that the species serves as an oracle, foretelling misfortunes for the village; its role as a symbol of grandparent identity; and the idea that colobus individuals do not intrude into croplands (Djègo-Djossou et al., 2012). Despite these protective traditional beliefs, the local community has witnessed a significant decline in the colobus population (Djègo-Djossou et al., 2012), largely due to hunting, particularly when the Kikélé monkeys venture into the unprotected adjacent gallery forests (Ota, 2020). It was also reported that the colobus population descended from a single pair of *C. vellerosus* brought from the village of Adjè in Togo to the Kikélé region around 1800 by Tchabi Mouin, a respected elder of the Tchabi-Ota family (Djègo-Djossou et al., 2012). Although the white-thighed colobus is held sacred in Kikélé and is protected by the local community, the KSF is degraded due to the unsustainable exploitation of the forest, the use of the forest as a toilet, and the deposit of garbage (Ota, 2020), and is fragmented due to agricultural activities and roads (Djègo-Djossou et al., 2015).

Given the assumed historical origin of only one pair of founder animals and the fragmentation of the KSF, inbreeding might pose an additional risk to the survival of this population. We therefore carried out an initial analysis of genetic diversity using mitochondrial markers, including the complete *cytochrome b* gene and a fragment of the hypervariable control region.

## Methods

### Sample collection

We collected 16 faecal samples non-invasively from two groups in the KSF and one group in the Okuta Kobunan Forest (OKF) between March 16, 2024, and April 2, 2024. Both forests are located in the Kikélé village (Fig. 2). In KSF, we monitored each group from dawn at 6:30 AM to 11:45 AM and again in the afternoon from 3:30 PM to 7:30 PM. During our observations, we collected fresh samples as soon as individuals defecated. Based on the characteristics of individuals, we cared not to sample the same individual twice. However, we could not completely rule this out. In OKF, we collected faecal samples from eight confidentially identified individuals of one group. Based on their size, samples were collected in either 15 ml or 50 ml Falcon tubes, stored in 96% ethanol, and transported from the field to the Laboratory of Applied Ecology of the University of Abomey-Calavi, Benin. There, they were preserved using silica gel, following the two-step protocol outlined by Nsubuga et al. (2004) and Roeder et al. (2004), until their transportation to the German Primate Center, Göttingen, Germany, for genetic analysis. For each sample, we recorded date, time, and geographic coordinates (Global Positioning System).

**Fig. 2:**
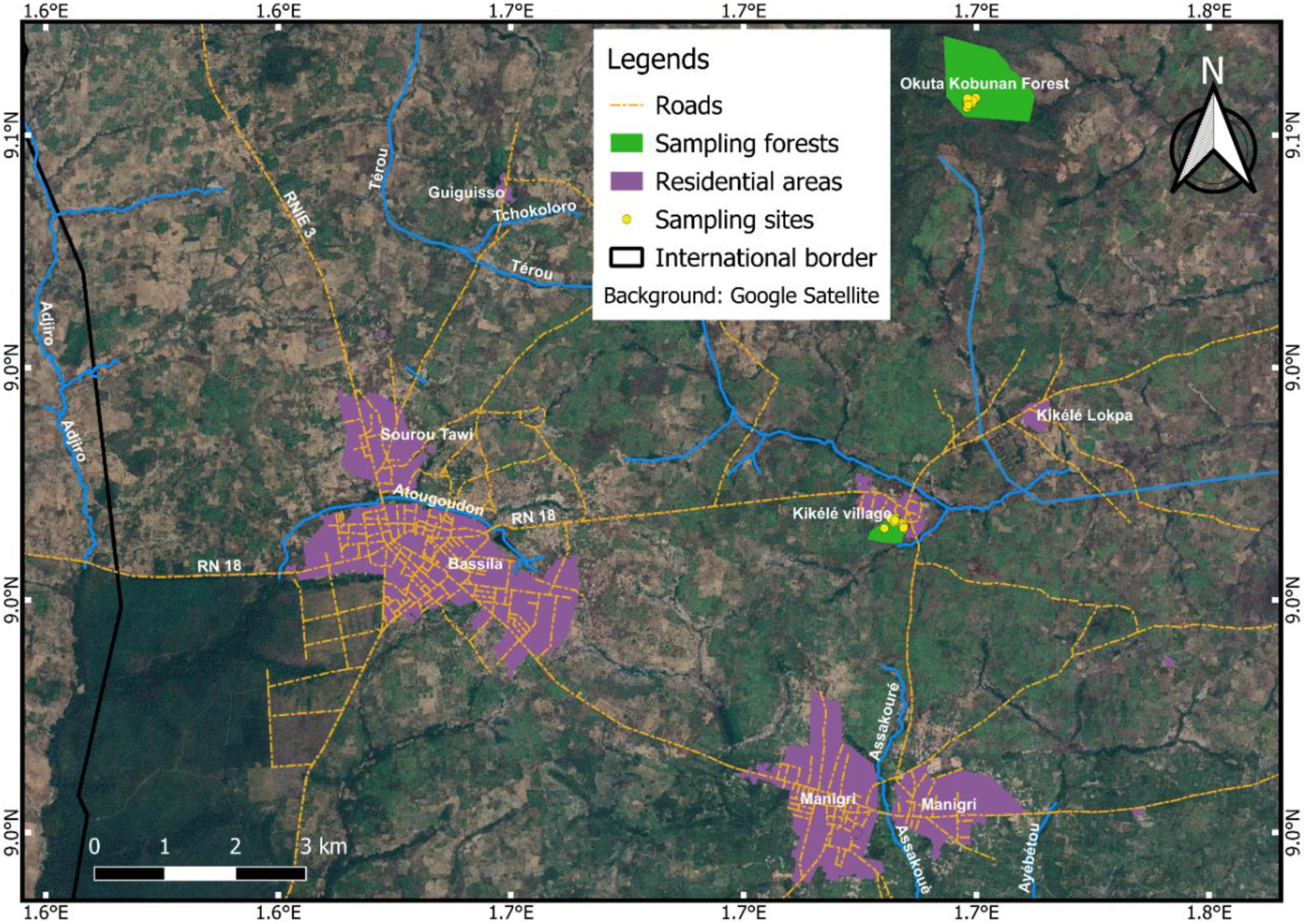
Sampling area around Kikélé village in west-central Benin (Kikélé Sacred Forest, KSF; Okuta Kobunan Forest, OKF).

### Laboratory work

We extracted total genomic DNA using the NucleoSpin® DNA Stool kit (Machery-Nagel), following the manufacturer’s protocols. After extraction, DNA concentrations (17-110 ng/μL) were quantified with a NanoDrop ND-1000 spectrophotometer (Peqlab). We stored the eluted samples at −20 °C until further analysis.

We amplified and sequenced two mitochondrial fragments, the complete cytochrome b gene (cytb; 1140 bp) and a fragment of the hypervariable control region (D-loop). We amplified cytb via two overlapping PCR products with sizes of 727 bp and 663 bp, while D-loop was amplified via a single PCR product with a size of 750 bp (for primers see Table 1).

**Table 1.**
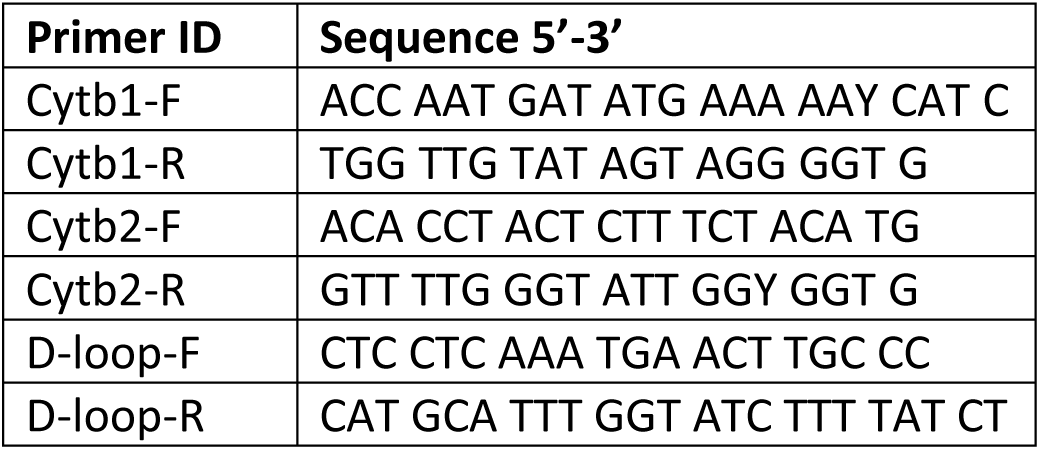
Primers used in this study

We performed polymerase chain reaction (PCR) in a total volume of 25 µl containing 2.5 µl 10X buffer, 200 µM dNTPs, 0.4 µM of each primer, 1 Unit FastStart Taq DNA polymerase (Roche), double-distilled water (ddH2O), and 50 ng template DNA. Each PCR run included an initial denaturation step at 95°C for 4 minutes, followed by 40 cycles of denaturation at 95°C for 30 seconds, annealing at 54°C (both cytb fragments) or 58°C (D-loop) for 30 seconds, and extension at 72°C for 60 seconds, concluding with a final extension at 72°C for 7 minutes, and subsequent cooling at 8°C. We ran the PCR product in 1% agarose gels stained with ethidium bromide (0.5 µg/mL) to check for PCR performance. We cut PCR products from the gel, cleaned them with the Monarch® DNA Gel Extraction Kit (New England Biolabs), and then sent them to Eurofins Germany for Sanger sequencing.

## Data analysis

We checked sequence electropherograms by eye and assembled and manually edited them in BioEdit Sequence Alignment Editor. We aligned the cytb and D-loop sequences using MUSCLE (Edgar, 2004) as implemented in MEGA 12 (Kumar et al., 2024). We also used MEGA12 to check for the correct translation of the cytb sequences into amino acid sequences.

To confirm species identity and marker specificity, we employed BLAST (NCBI). However, since no orthologous cytb sequences of *C. vellerosus*, *C. polykomos,* and *C. satanas* are available in GenBank, we used complete cytb sequences of *C. guereza* and *C. angolensis* to determine the relationship of *C. vellerosus* to these two other *Colobus* species. In a second step, we downloaded orthologous D-loop sequences from *C. polykomos*, *C. guereza,* and *C. angolensis*. We used *Piliocolobus badius* as an outgroup and reconstructed the phylogenetic trees for cytb and D-loop using the Maximum Likelihood (ML) algorithm with 100 bootstrap replications in MEGA12.

Utilising the aligned cytb and D-loop sequences, we assessed the genetic diversity within the population of *C. vellerosus* inhabiting the KSF and OKF by calculating the number of variable sites (s), total number of mutations (η), haplotype diversity (Hd), and nucleotide diversity (π) with the DnaSP software v6.12 (Rozas et al., 2017).

## Results

We successfully generated complete cytb and partial D-loop sequences from four and fifteen samples, respectively. Among them, we found only one haplotype for cytb and two haplotypes for D-loop.

As expected, the ML tree (Fig. 3) showed that the cytb haplotype from the *C. vellerosus* population in the Kikélé village forests forms a sister lineage to *C. guereza*, albeit with low statistical support (79%). The D-loop sequences form a sister clade to *C. polykomos* from Guinea-Bissau, but also with low bootstrap support of only 64%. *Colobus guereza* is sister to the *C. vellerosus* + *C. polykomos* clade, while *C. angolensis* represents to most basal species among the investigated *Colobus* species.

**Fig. 3:**
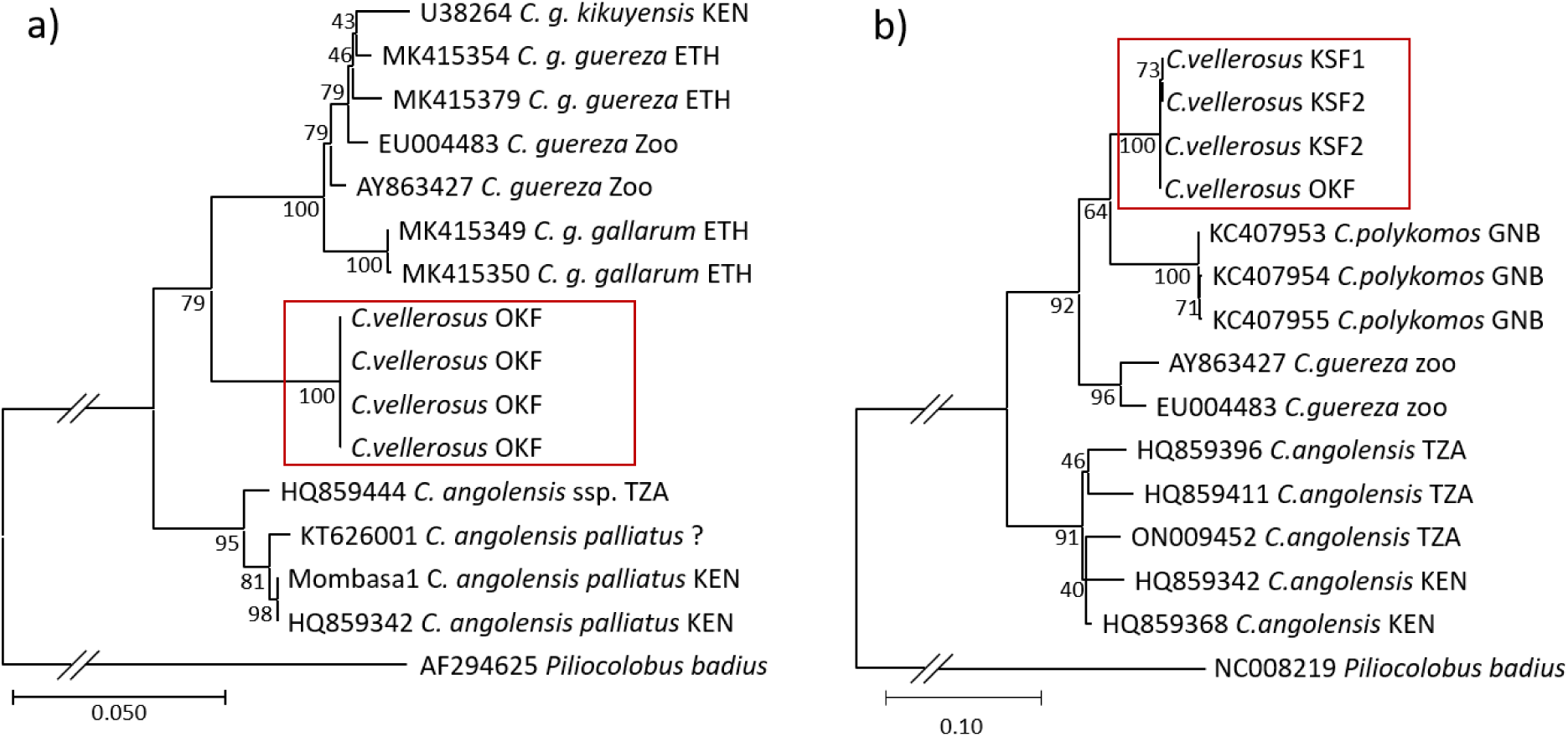
Phylogenetic relationships of *Colobus vellerosus* based on a) the complete cytochrome b gene 1140 bp and b) a 484 bp long fragment of the D-loop. Orthologue sequences were downloaded from GenBank, *Piliocolobus* was used as an outgroup. The phylogeny was inferred using the Maximum Likelihood method and the HKY+G+I model (Hasegawa et al., 1985). The percentage of replicate trees in which the associated taxa clustered together (100 bootstrap replicates) is shown next to the branches. For the D-loop, we used shorter sequences than 750 bp due to the shorter lengths of the orthologue sequences from GenBank. The bar below the trees indicates substitutions per site.

The two D-loop haplotypes differ only at one position. One haplotype was found in three individuals of the group from the OKF, and one of the second group (group 2) from the KSF. The second haplotype was found in six individuals from the first group (group 1) of the KSF and five individuals from group 2 of the KSF. The estimated haplotype diversity (Hd) was 0.419 ± 0.113, while nucleotide diversity (π) was 0.00056 ± 0.00015, indicating very low genetic diversity in the mitochondrial non-coding region.

## Discussion

We found no genetic variability in the four cytb sequences and only minimal polymorphism in the D-loop within the *Colobus vellerosus* population around Kikélé village. This low mitochondrial genetic diversity suggests a potential recent genetic bottleneck, founder effect, or sustained genetic drift, particularly relevant for small or isolated populations (Ellegren & Galtier, 2016; Mayr, 1966; Nei et al., 1975).

Historically, the Kikélé colobus population is believed to originate from a single pair of *C. vellerosus* (Djègo-Djossou et al., 2012). Given that mitochondrial DNA is inherited through the maternal lineage (Giles et al., 1980; Gutierrez et al., 2024), the monomorphic mitochondrial DNA indicates a common maternal ancestor among individuals, supporting the founder effect hypothesis.

The forests adjacent to the Kikélé village, including the Pénéssoulou Forest Reserve, Wari-Maro, Ouémé Supérieur, and Monts Kouffé, served as habitats for *C. vellerosus* (Djègo-Djossou & Sinsin, 2009), most likely harbouring additional mitochondrial haplotypes. These haplotypes have most likely gone lost when *C. vellerosus* became extinct, due to the degradation of these forests and hunting (Djègo-Djossou & Sinsin, 2009; Orekan, 2007; Ota, 2020). The fur of *C. vellerosus* was used in the production of local shoes and drums (Gautier-Hion et al., 1999; Oates, 2011), and their bones were utilised in traditional medicine and magic (Djagoun et al., 2018).

Another consideration is that our current sampling may not have captured all haplotypes present within the Kikélé population. Nonetheless, relying solely on mitochondrial DNA analysis is insufficient to conclude about the genetic diversity of wild animal populations (Lynn et al., 2016; Roos & Zinner, 2017).

### Mitochondrial genetic diversity

In our study, we identified two haplotypes that differ at a single site within the mitochondrial D-loop. Similarly, Ting (2008) found only a single mitochondrial haplotype in four samples from the *C. vellerosus* population at the Boabeng Fiema Monkey Sanctuary in Ghana. If this locally low mitochondrial genetic diversity is a general characteristic of the species, its conservation status might be even worth than reported.

However, the low mitochondrial genetic variability within local populations of *C. vellerosus* seems not to be an exception among colobines. Several populations of Asian colobines show very low to no mitochondrial genetic variability. Ang et al. (2012) reported only two haplotypes in the Raffles’ banded langur (*Presbytis femoralis*) population in Singapore, differing by a single nucleotide in the D-loop. In 201 samples of the largest population of the Tonkin snub-nosed monkey (*Rhinopithecus avunculus*), Ang et al. (2016) did not find any mitochondrial genetic variability. Similarly, in 128 samples of the grey or Guizhou snub-nosed monkey (*Rhinopithecus brelichi*), only one mitochondrial haplotype was found (Pan et al., 2011). However, in contrast to the low mitochondrial genetic variability, in this species, the nuclear genetic diversity, as determined by allele numbers of microsatellites, was significantly higher (Kolleck et al., 2013). The contrast might be the result of interspecific gene flow between *R. roxellana* and the ancestor of *R. bieti* and *R. strykeri* (Wu et al., 2023). That most likely led to hybrid speciation where males of only one lineage introgressed the population of the other lineage and thus importing nuclear variability but not mitochondrial variability. To obtain a complete picture of the genetic diversity in these endangered species, population genetic analyses based on nuclear markers are needed.

Within other species of the genus *Colobus*, McDonald et al. (2022) detected a minimum of one mitochondrial haplotype in two distinct populations and a maximum of seven haplotypes in one population from 140 sequenced samples collected from nine *C. angolensis* populations in southern Kenya and the Southern Highlands of Tanzania. In total, 26 haplotypes were identified across these nine studied populations (McDonald et al., 2022; McDonald & Hamilton, 2010). Although details regarding mitochondrial genetic diversity at the population level in *C. g. guereza* and *C. g. gallarum* are limited; Zinner et al. (2019) listed 76 sequences, including 31 haplotypes for cytb from samples collected across Ethiopia. For *C. polykomos*, three different mitochondrial haplotypes were detected in the Cantanhez Forests National Park in Guinea-Bissau (Minhós et al., 2013), while five different haplotypes were found in the Taï National Park in Ivory Coast (Minhós et al., 2023). Currently, no genetic data is available for *C. satanas*.

### Implications for conservation

Low genetic diversity presents challenges to the sustainability of populations, particularly under environmental stressors or disease threats. Within different animal taxa, ensuring adaptability and disease resilience hinges on fostering genetic variation (Ostridge et al., 2025; Schwensow et al., 2007; Tallmon et al., 2004; Warren et al., 2025; Zhang et al., 2024). To address this issue, comprehensive conservation strategies must be employed that encompass several key components. Firstly, improving habitat connectivity is essential to facilitate gene flow between populations, thereby minimising genetic isolation. This can be achieved through the creation and maintenance of ecological corridors that enable individual movement across fragmented landscapes. Regular population monitoring is vital for understanding population dynamics over time, allowing for the identification of potential demographic decline. Moreover, a comprehensive study on the global genetic diversity utilising both mitochondrial and nuclear markers should be carried out. This approach will provide a detailed understanding of the genetic structure of the species. Establishing a localised genetic database would greatly enhance conservation efforts, serving as a critical repository for genetic information that can inform management strategies, such as defining management units, genetic rescue by outcrossing or translocation, reintroduction, and connectivity network for meta-population. In instances where populations exhibit severe genetic isolation and decline, translocation may be necessary as a genetic rescue strategy. Relocating individuals from genetically not too diverse donor populations into the vulnerable groups can augment their gene pool, thereby improving their reproductive viability and bolstering resilience against extinction pressures. Such proactive measures are essential to ensure the longevity of threatened populations under the current rapidly changing anthropogenic and climate change environment.

## Conclusion

Our research contributes to the dire conservation status of *C. vellerosus*, demonstrating that not only is the global population size small, but also the genetic diversity is most likely extremely low. Our study also indicates that a range-wide population genetic study is needed to explore the possibilities for genetic rescue by exchanging individuals among local populations.

## Data availability

Sequence data from this study are available from NCBI GenBank: cytb PV963062 - PV963065; D-loop PV963066 - PV963080.

## Author contributions

Conceptualisation: FDA, CAMSD, CR, and DZ; data collection: FDA and DD; laboratory work: FDA and DJZ; data curation and formal analysis: FDA and DZ; writing-original draft: FDA and DZ; writing-reviewing and editing: FDA, SSMT, IK, DJZ, DD, CAMSD, CR, and DZ; supervision and resources: SSMT, IK, CR, and DZ.

## Acknowledgments

The authors express their gratitude to the German Federal Ministry of Education and Research (BMBF) and the West African Science Service Centre on Climate Change and Adapted Land Use (WASCAL) for funding this study through the Climate Change and Biodiversity Doctorate Research Program at Université Félix Houphouët-Boigny (UFHB) in Côte d’Ivoire. We also thank re:wild and the Förderkreis of the German Primate Center for financial support. We extend our heartfelt thanks to our field guides, Mr. Ganiou Adjofè and Mr. Dramani Gado, who assisted us during data collection in Kikélé village in central Benin. Sadly, Mr. Dramani Gado passed away before we completed this work; may his soul rest in peace.

## Ethical Statement

This research protocol was approved by the Research Committee of Wildlife Conservation at the Laboratory of Applied Ecology, University of Abomey-Calavi, Benin. We used non-invasive faecal sampling methods that comply with Benin’s laws and the guidelines established by the International Primatological Society.

